# A Simultaneous Inhibition of ID1 and ID3 Protects Against Pulmonary Fibrosis

**DOI:** 10.1101/2025.07.24.665373

**Authors:** Samar A Antar, Eric Mensah, Jacob Dahlka, Michael Aziz, Aymen Halouani, Seun Imani, Aanandi Parashar, Ahmed A Raslan, Robert Benezra, Diego Fraidenraich, Giovanni Ligresti, Yassine Sassi

## Abstract

**Background:** Idiopathic pulmonary fibrosis (IPF) is a fatal lung disease for which novel therapeutic approaches are desperately needed. Inhibitor of DNA binding (ID) proteins are regulated by Transforming Growth Factor-β. However, the regulation and the effects of ID proteins in IPF remain poorly understood. We aimed to assess the expression of ID proteins in IPF and determine the effects of ID proteins on human lung fibroblasts (HLF) *in vitro* and pulmonary fibrosis *in vivo*.

**Methods:** The expression of ID proteins in lungs and lung fibroblasts from mice and human patients with pulmonary fibrosis was evaluated. The effects of ID1/ID3 inhibition and overexpression on HLF were assessed. Genetic and pharmacological approaches were used *in vivo* to determine the role of ID1/ID3 in pulmonary fibrosis.

**Results:** ID1/ID3 levels were elevated in HLFs isolated from pulmonary fibrosis-diseased patients and mice. ID1/ID3 knockdown decreased IPF-diseased HLF proliferation and differentiation into myofibroblasts. Bleomycin-exposed ID1/ID3 KO mice displayed improved lung function and presented with decreased lung fibrosis when compared to WT mice. A pharmacological inhibitor of ID1/ID3 decreased IPF-diseased HLF proliferation and differentiation *in vitro* and attenuated pulmonary fibrosis *in vivo*. A lung specific inhibition of ID1/ID3, using adeno-associated viruses expressing short hairpins targeting ID1 and ID3, reversed pulmonary fibrosis in mice. Mechanistically, ID1/ID3 inhibition decreased fibroblast proliferation through cell cycle genes and inhibited fibroblast differentiation through the MEK/ERK pathway.

**Conclusions:** Our data indicate that a simultaneous inhibition of ID1 and ID3 attenuates pulmonary fibrosis. ID1/ID3 inhibition holds potential as a novel therapeutic treatment for IPF.

## Introduction

Idiopathic pulmonary fibrosis is (IPF) is an irreversible, chronic, progressive fibrosing lung disease of unknown cause with a median survival time of 3-5 years post-diagnosis (1, 2). IPF affects older adults and is characterized by a gradual decline in lung function and worsening dyspnea, resulting in a poor prognosis. The exact causes and mechanisms of lung fibrosis remain unclear. However, it is well known that environmental, age-related, and genetic factors contribute directly to alveolar epithelial injury (3). It is currently thought that the persistent activation of fibroblasts leads to the accumulation of extracellular matrix (ECM) in the interstitium, which in turn reduces alveolar spaces (4). Current medications merely slow down disease progression but are insufficient to completely halt it. Therefore, new therapeutic approaches are desperately needed.

Inhibitors of DNA Binding (ID1-4) proteins are a subgroup of the helix-loop-helix (HLH) transcription factor family, which play a crucial role in regulating cell fate (5). However, unlike other HLH members, they lack a DNA-binding motif. ID proteins function as a dominant negative regulator of other HLH proteins by inhibiting their DNA binding and transcriptional activity (5). These proteins are significant in development, where they regulate processes such as cell-cycle progression, cell proliferation, differentiation, cell fate determination, hematopoiesis, angiogenesis, and the metabolic adaptation of cancer cells (6–9). However, the effects of ID proteins in IPF pathogenesis remain poorly understood. Addressing these knowledge gaps by elucidating the role of ID proteins in IPF pathogenesis and progression could reveal novel therapeutic avenues, directly responding to the urgent need for more effective IPF treatments.

In the present study, we investigated the regulation and role of ID proteins in pulmonary fibrosis using a combination of genetic and pharmacological approaches. Our study demonstrates that inhibiting simultaneously ID1 and ID3 attenuates the proliferation, migration, and differentiation of human lung fibroblasts *in vitro*, and the development of pulmonary fibrosis *in vivo* in mice.

## Materials and Methods

Details of the materials and can be found in the supplementary material.

### Human lung fibroblasts

Normal (healthy) human lung fibroblasts (NHL-FBs; cat #CC-2512) isolated from 6 healthy donors, and IPF-diseased human lung fibroblasts (IPF-FBs; cat#CC-7231) isolated from 6 patients with IPF were purchased from Lonza and cultured according to the recommended guidelines. The cells were passed upon reaching confluency and were used at passages 2 to 7. All cells used in this study were tested to confirm the absence of HIV-1, HBV, HCV, mycoplasma, bacteria, yeast and fungi.

### Intratracheal BLM mouse model

All the animal experiments were performed in accordance with NIH Guide for the Care and Use of Laboratory Animals. Approval was obtained from Institutional Animal Care at Mount Sinai and Virginia Tech. C57BL/6 male mice were anesthetized with ketamine/xylazine before being immobilized in the supine position on the operating field, and intubated using a 20G catheter. The BLM solution (2 units/kg) was delivered by direct injection into the trachea using a syringe. Once delivery was complete, the needle was withdrawn, and the animals were extubated and placed in a warm environment until they fully recovered from anesthesia, after which they were returned to their respective cages.

### AAV delivery

The mice were anesthetized by IP injection of xylazine and ketamine and were secured to a tray in the supine position. Subsequently, using a 20G angiocath, animals were intubated. The board was tilted at 45 degrees and the IA-1C Micro sprayer tip (PennCentury) was inserted through the lumen of the angiocath. AAV1-shRNA-ID1/ID3 (Vector Biolabs) and AAV1-Luciferase (AAV-Ctrl) were administered via IT delivery (3x10^11^ in 100 μL). The animals were then extubated and returned to their cages.

### Statistical analyses

All quantitative data are reported as means ± SEM. Statistical analysis was performed with the Prism software package (GraphPad Version 10). Differences between two means were assessed by a two-tailed paired or unpaired t-test. Differences among multiple means were assessed by one-way or two-way ANOVA followed by Bonferroni correction or followed by Holm–Sidak’s test analysis. P-values <0.05 were considered significant (corresponding symbols in figures are *for P < 0.05, **for P < 0.01, and ***for P < 0.001).

## RESULTS

### ID1 and ID3 are upregulated in pulmonary fibrosis

In order to investigate the regulation of ID family members in pulmonary fibrosis, we first used publicly available single cell RNA sequencing data set from Healthy and IPF lungs (GSE132771) (10). We found ID1 and ID3 levels to be significantly increased in alveolar fibroblasts from IPF patients, which are the primary source of CTHRC1+ pathogenic collagen-producing fibroblasts (11) (figure 1a). We next quantified ID1 and ID3 mRNA levels in human lung fibroblasts (HLF) isolated from healthy donors and from patients with IPF. The PCR analyses revealed a significant increase in ID1 and ID3 levels in diseased fibroblasts (figure 1b). Human lung fibroblasts were then treated with TGF-β, the primary driver of pulmonary fibrosis (12), and ID1 and ID3 mRNA levels were quantified by quantitative PCR. Our analyses revealed a prominent increase of ID1 and ID3 mRNA levels in TGF-β-treated fibroblasts (figure 1c). Next, we explored ID1/ID3 mRNA expression profiles in an *in vivo* animal model of lung fibrosis: Bleomycin (BLM)-induced pulmonary fibrosis in mice. The PCR analyses revealed a significant increase in pulmonary ID1 and ID3 mRNA levels in diseased mice (figure 1d). Consistent with the qPCR results, we found pulmonary ID1 and ID3 protein levels to be increased in the lungs of mice 3 weeks after BLM delivery (figure 1e). Additionally, we examined the mRNA expression profiles of ID1 and ID3 in fibroblasts isolated from the lungs of mice with pulmonary fibrosis. PCR and immunoblotting analyses revealed a significant increase in ID1 and ID3 mRNA and protein levels in fibroblasts isolated from the lungs of diseased mice (figure 1f-g). Importantly, no significant changes in ID2 and ID4 levels were observed in lung fibroblasts from IPF patients, healthy lung fibroblasts treated with TGF-β or lungs of mice with pulmonary fibrosis (supplementary figure S1). These results indicate that ID1 and ID3 levels are increased in lung fibroblasts from mice and humans with pulmonary fibrosis, suggesting a contribution of these proteins to disease progression.

**Figure 1:**
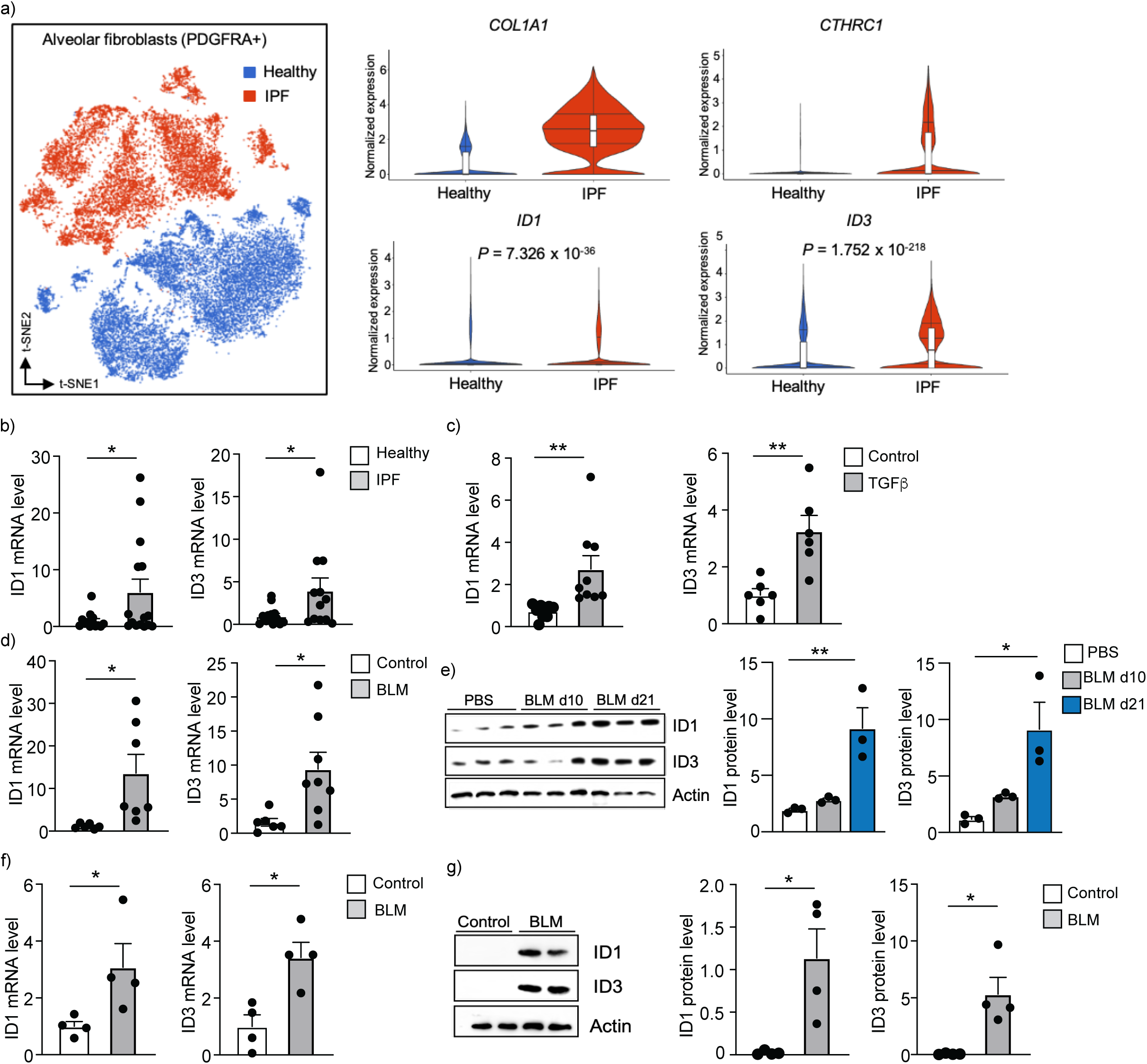
ID1 and ID3 regulation in pulmonary fibrosis. a) (Left) t-SNE plot displaying alveolar fibroblasts Pdgfra+ clusters in healthy (blue) and IPF (red) lungs. (Right) Violin plots showing the expression of Col1a1, Cthrc1, ID1 and ID3 genes in healthy and IPF-diseased lungs. b) ID mRNA levels determined by real time qPCR analyses in lung fibroblasts isolated from healthy donors and patients with IPF (n = 12-15/group). c) ID mRNA levels in healthy human lung fibroblasts treated with TGF-β1 (5 ng/ml) for 48 hours. n = 6-9 experiments performed in triplicate. d) ID1 and ID3 mRNA levels in lungs of mice with pulmonary fibrosis (n = 6-8 mice/group). e) ID1 and ID3 protein levels in lung homogenates 10 and 21 days after PBS or BLM injection (n = 3 mice/group). f) ID1 and ID3 mRNA levels in lung fibroblasts isolated from the lungs of PBS or BLM-treated mice (n = 4 mice/group). g) ID1 and ID3 protein levels in lung fibroblasts from control and BLM-treated mice (n = 4 mice/group). * P < 0.05, ** P < 0.01.

### ID1 and ID3 regulate human lung fibroblast proliferation and differentiation into myofibroblasts

Given that lung fibroblasts contribute to injury responses through rapid migration, proliferation, and differentiation into myofibroblasts, we assessed whether ID1/ID3 inhibition affects these mechanisms in HLF. ID1 knockdown using a specific siRNA induced a compensation by increased ID3 level, and increased ID1 level compensated ID3 knockdown (figure 2a), indicating that a simultaneous inhibition of ID1 and ID3 is essential to characterize their roles in pulmonary fibrosis. A simultaneous knockdown of ID1/ID3 significantly prevented healthy and IPF-diseased HLF proliferation induced by serum (5% FBS; figure 2b). We next assessed whether ID1/ID3 inhibition affects HLF migration. Serum treatment induced an increase in healthy and IPF-diseased lung fibroblast migration, whereas ID1/ID3 knockdown significantly prevented this effect (figure 2c-d).

**Figure 2:**
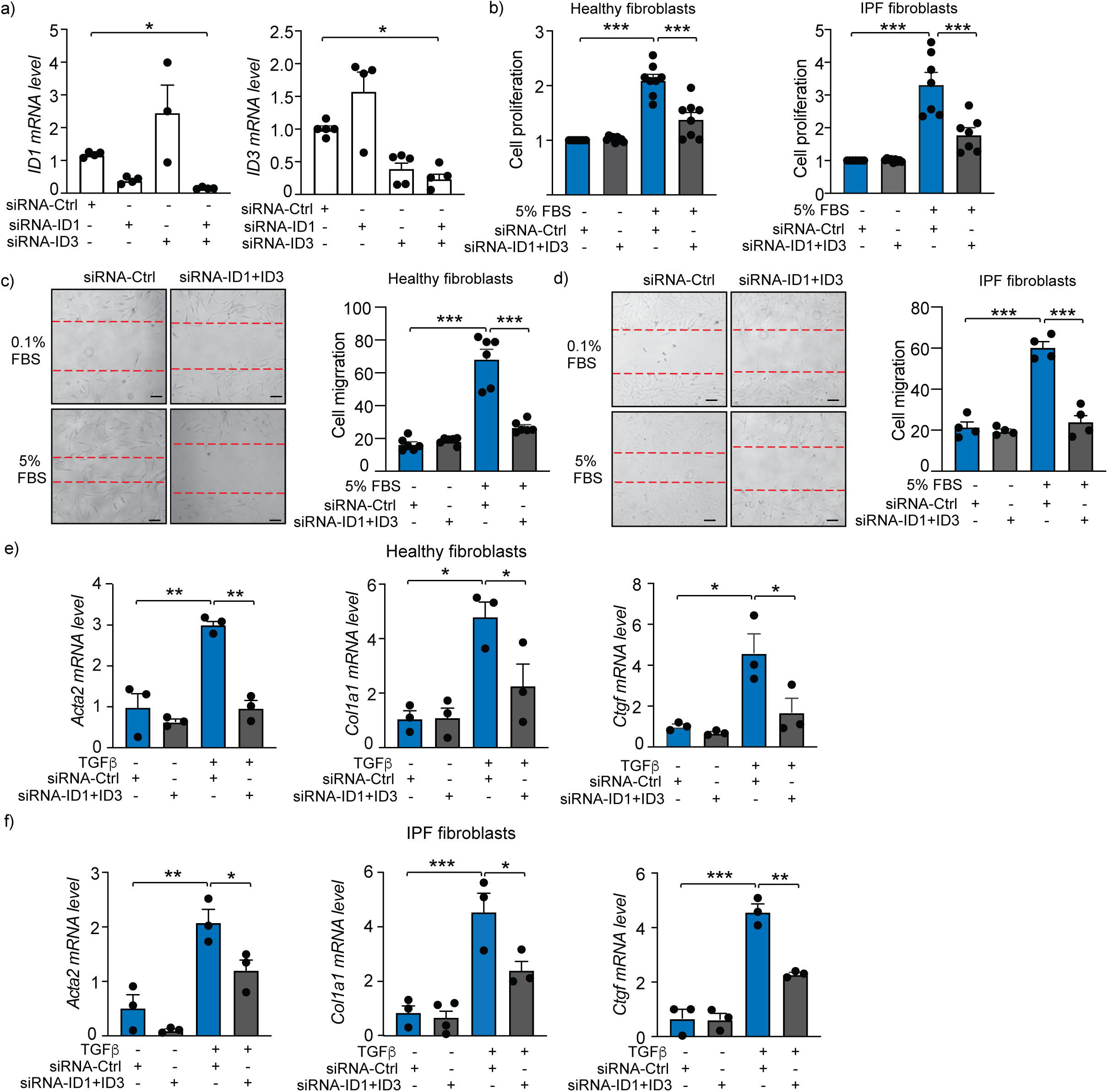
**Knockdown of ID1/ID3 decreases human lung fibroblast proliferation, migration and differentiation into myofibroblasts**. a) ID1 and ID3 mRNA levels in healthy human lung fibroblasts transfected with siRNA-Ctrl, siRNA-ID1 or siRNA-ID3 (5nM each) for 48 hours. n = 3-5 experiments performed in triplicate. b) Proliferation of healthy and IPF-diseased human lung fibroblasts in the presence of the indicated treatments. n = 7-9 experiments performed in triplicate. (c-d) Migration of healthy (c) and IPF-diseased (d) human lung fibroblasts in the presence of the indicated treatments. n = 4-6 experiments performed in duplicate. Scale bar: 100 µm. (e-f) qPCR assessment of Acta2, Col1a1 and CTGF mRNA levels 48h in healthy (e) and IPF-diseased (f) HLF treated with siRNA-Ctrl or siRNA-ID1ID3 in the absence or presence of TGF-β1 (5 ng/ml). n = 3-4 experiments performed in triplicate. * P < 0.05, ** P < 0.01. *** P < 0.001.

Since fibroblasts differentiate into myofibroblasts to increase collagen production and promote fibrosis, we then assessed whether ID1/ID3 inhibition can affect this process. Healthy and IPF-diseased HLF were treated with TGF-β1 in the presence or absence of ID1 and ID3 specific siRNAs. As an indicator of fibroblasts differentiation, TGF-β1 increased alpha smooth muscle actin (Acta2) mRNA level, as well as Collagen 1a1 (Col1a1) and Connective tissue growth factor (Ctgf) mRNA levels (figure 2e-f). Importantly, ID1/ID3 knockdown prevented these changes (figure 2e-f), thus indicating that ID1/ID3 inhibition impairs lung fibroblast differentiation. These results indicate that ID1/ID3 specific knockdown decreases lung fibroblast proliferation, migration and differentiation into myofibroblast.

We next investigated the effects of ID1/ID3 overexpression on HLF proliferation, migration and differentiation into myofibroblast. HLF, isolated from human healthy donors, were treated with adenoviruses ID1 and ID3 (Ad-ID1+Ad-ID3) or with an adenovirus encoding Luciferase (Ad-Ctrl, as a control). A significant increase in ID1 and ID3 levels was observed in Ad-ID1/ID3-treated HLF (supplementary figure S2a), showing the efficiency of the generated adenoviruses. ID1/ID3 overexpression did not affect healthy HLF proliferation but induced a significant increase in fibroblast migration (supplementary figure S2b-c). Importantly, ID1/ID3 overexpression induced Acta2, Col1a1 and Ctgf mRNA levels (supplementary figure S2d) to levels comparable to those observed upon TGFβ1 treatment (figure 2). These results suggest that ID1/ID3 overexpression is sufficient to increase human lung fibroblast migration, to induce fibroblast differentiation into myofibroblast and to increase collagen production.

### ID1/ID3 KO mice are protected from BLM-induced pulmonary fibrosis

To evaluate the *in vivo* effect of ID1/ID3 deletion on pathological remodeling in lung fibrosis, we generated ID1/ID3 knock-out mice. Given that ID1/ID3 double knockout mice die at mid-gestation, we ablated the ID3 gene and conditionally ablated the ID1 gene in fibroblasts to generate conditional KO mice. These mice were developed by crossing mice carrying a ubiquitous deletion of ID3 with mice carrying a fibroblast specific deletion of ID1 (by crossing ID1^fl/fl^ mice with Col1a2-CreER mice). Mice received a daily injection of tamoxifen via IP administration during 5 days (figure 3a). Mice were then intratracheally injected with a single dose of BLM (2U/kg), or a saline solution and were sacrificed four weeks after BLM delivery (figure 3a). As expected, BLM treatment in WT mice resulted in a marked increase in ID1 and ID3 levels, whereas ID1/ID3 KO mice displayed a marked decrease in pulmonary ID1 and ID3 mRNA levels (figure 3b). Importantly, no significant changes in ID2 and ID4 levels were observed in the lungs of the ID1/ID3 KO mice (supplementary figure S3a-b). Disease progression in patients with IPF is often assessed using imaging and lung function analyses (13); and in clinical studies, improvements in lung function are primarily used to gauge the success of therapeutics directed towards IPF. Therefore, we performed lung function analyses of the used mice. BLM-exposed ID1/ID3 KO mice showed significant improvements in all measured parameters that were altered in BLM-treated WT mice, including increased inspiratory capacity, static compliance, and decreased respiratory elastance (figure 3c). Immunoblot analyses of Collagen-I and α-SMA expression in lung homogenates showed increased Collagen-I and α-SMA protein levels in response to BLM in WT mice, whereas ID1/ID3 KO mice displayed a significant decrease in their expression levels (figure 3d). Similarly, we found Collagen 3a1 (Col3a1) and Ctgf levels to be increased in response to BLM in WT mice, whereas ID1/ID3 KO mice displayed a significant decreased in their mRNA levels (supplementary figure S3c-d). Histological examination of lung tissues by Sirius Red/Fast Green staining showed a fibrotic reaction in response to BLM in WT mice, whereas ID1/ID3 KO mice displayed a decrease in lung fibrosis (figure 3e). Consistently, in comparison to the WT group, ID1/ID3 KO mice presented with a lower Ashcroft score, reflecting decreased fibrotic changes (figure 3e). Finally, ID1/ID3 KO mice presented a significant decrease in hydroxyproline level when compared to BLM-treated WT mice (figure 3f). Taken together, these results indicate that deletion of ID1*/*ID3 prior to BLM exposure protects mice from the development of lung fibrosis.

**Figure 3:**
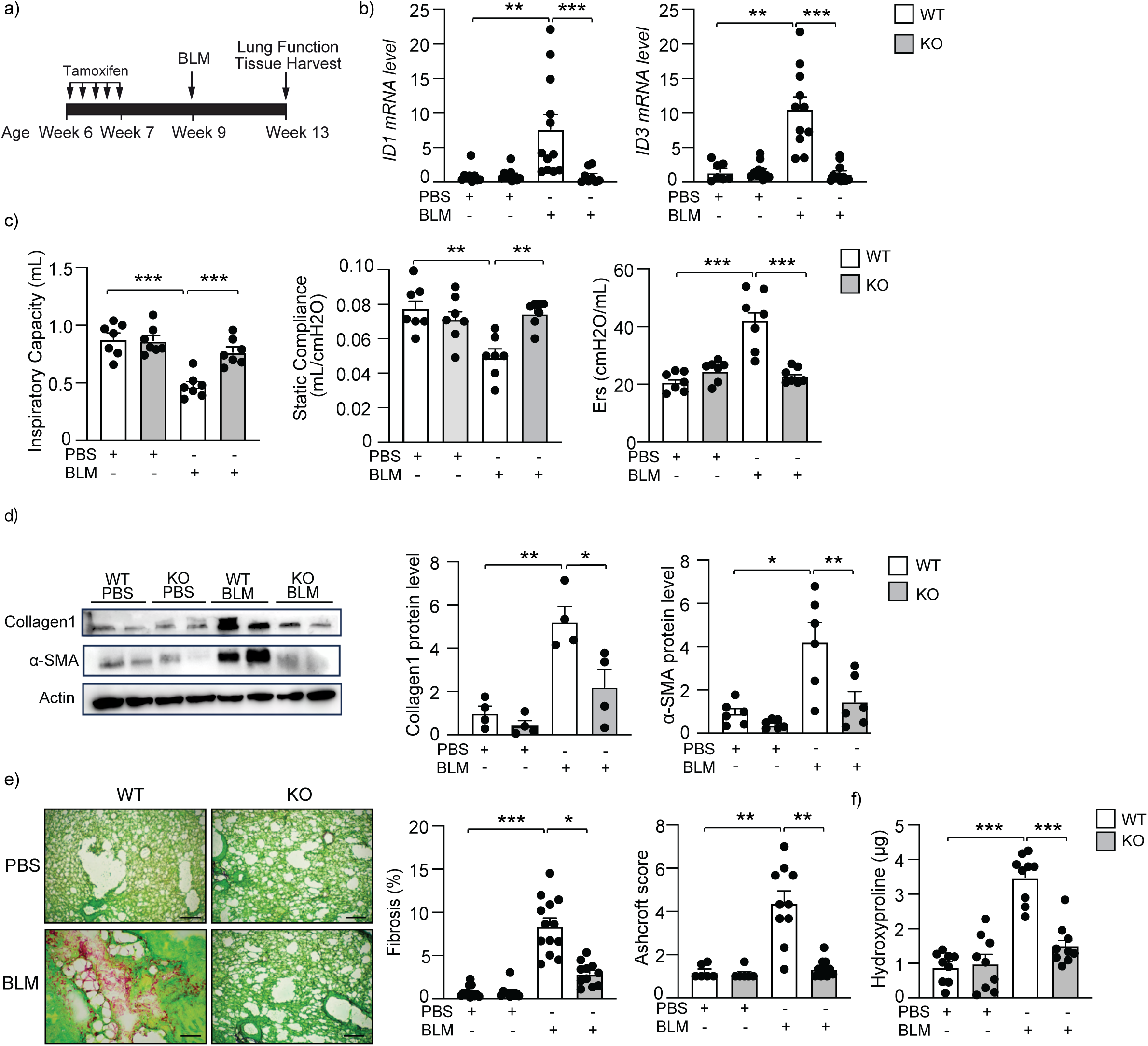
ID1/ID3 deletion decreases Bleomycin-induced lung fibrosis. a) Design of the study. b) PCR analysis of ID1 and ID3 mRNA levels in lungs of the indicated groups n=7-12 mice/group. c) Lung function data parameters for inspiratory capacity, compliance and single frequency elastase n=7 mice/group. d) Lung protein expression of Collagen1 and Acta2 in lungs from the indicated groups n=4-6 mice/group. e) (Left) Representative images from Fast Green/Sirius-Red- stained lungs of the indicated groups n=10-13mice/group. Scale bars: 200 μm. (Middle) Quantitative analysis of fibrosis. (Right) Ashcroft scores representing the extent of fibrosisn= 6- 10 mice/group. f) Hydroxyproline content in lungs from the indicated groups. n = 9 mice/group. * P < 0.05; ** P < 0.01. *** P < 0.001.

### Pharmacological inhibition of ID1/ID3 attenuates pulmonary fibrosis

We next assessed whether a specific pharmacological inhibitor of ID1 and ID3 (AGX51) affects human lung fibroblasts proliferation, migration and differentiation into myofibroblasts in vitro. This small molecule leads to ubiquitin-mediated degradation of ID1 and ID3 (14). We found AGX51 to inhibits healthy and IPF-diseased HLF proliferation induced by serum (figure 4a-b). Consistent with the results of the experiments performed using ID1 and ID3 siRNAs, we found AGX51 to inhibits healthy and IPF-diseased human lung fibroblasts migration (figure 4c-d). In addition, AGX51 reduced the differentiation of both healthy and IPF-diseased HLFs, as assessed by qPCR and immunoblot analyses of fibrotic markers (figure 4e-f and supplementary figure S4). To assess the in vivo effects of AGX51 on pulmonary fibrosis, WT mice were intratracheally injected with BLM (2U/kg) or a saline solution, followed by two weeks of treatment with AGX51(delivered via IP injection) starting at day 14 from bleomycin adminstration (figure 5a). Under basal conditions (i.e., in healthy mice), AGX51 treatment did not elicit a lung phenotype (figure 5b-f). AGX51 treatment improved the lung function of BLM-treated mice, as evidenced by increased respiratory capacity and static compliance, and reduced respiratory elastance (figure 5b). The expression levels of Col1a1, Col3a1, and fibronectin (*Fn1*) were significantly reduced in the lungs of pulmonary fibrosis-diseased mice that were treated with AGX51 (figure 5c). Immunoblot analysis of collagen-I and collagen-III protein levels in lung homogenates also showed a marked decrease in collagen production in response to AGX51 treatment (figure 5d). Histological examination of lung tissues through Sirius Red/Fast Green staining revealed a fibrotic response to BLM in WT mice, whereas AGX51 treatment led to a reduction in lung fibrosis (figure 5e). Similarly, AGX51-treated mice exhibited a lower Ashcroft score compared to the PBS-treated group, indicating reduced fibrotic changes (figure 5e). In addition, AGX51-treated mice showed a significant decrease in hydroxyproline level (figure 5f). Together, these results indicate that pharmacological inhibition of ID1 and ID3 protects mice from the development of lung fibrosis.

**Figure 4:**
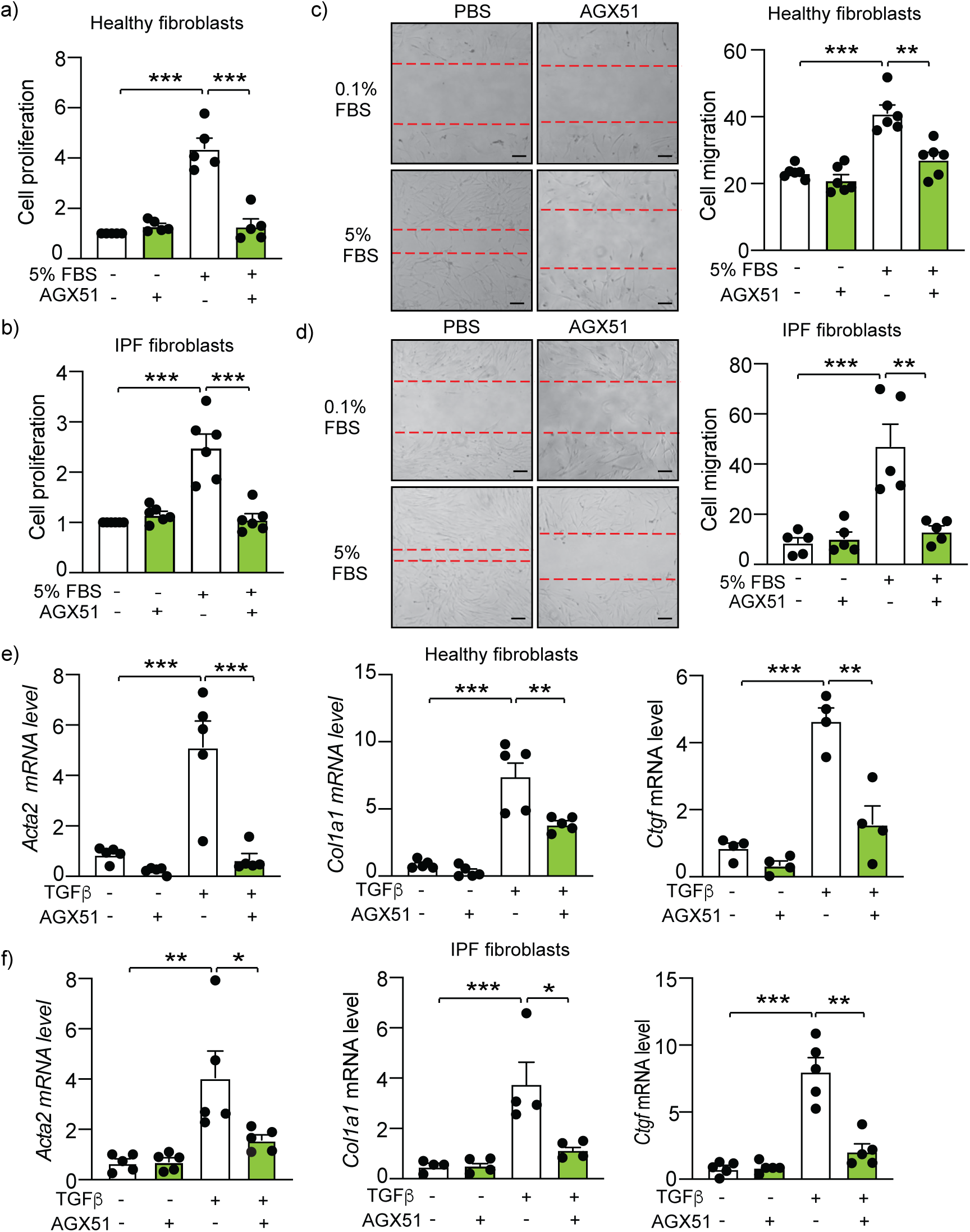
A pharmacological inhibition of ID1/ID3 reverses IPF-diseased HLF proliferation and differentiation into myofibroblasts. (a-b) Proliferation of healthy (a) and IPF-diseased (b) human lung fibroblasts in the presence or absence of an ID1/ID3 pharmacological inhibitor (AGX51, 20µM). n = 5 experiments performed in triplicate. (c-d) Migration of healthy (c) and IPF-diseased (d) human lung fibroblasts in the presence or absence of an ID1/ID3 pharmacological inhibitor (AGX51, 20µM). n = 5-6 experiments performed in triplicate. Scale bar: 100 µm. (e-f) qPCR assessment of Acta2, Col1a1 and Ctgf mRNA levels 48h after healthy (e) and IPF-diseased (f) human lung fibroblasts treatment with TGF-β1 (5 ng/ml) and AGX51 (20 µM). n = 4-5 experiments performed in triplicate. * P < 0.05; ** P < 0.01. *** P < 0.001.

**Figure 5:**
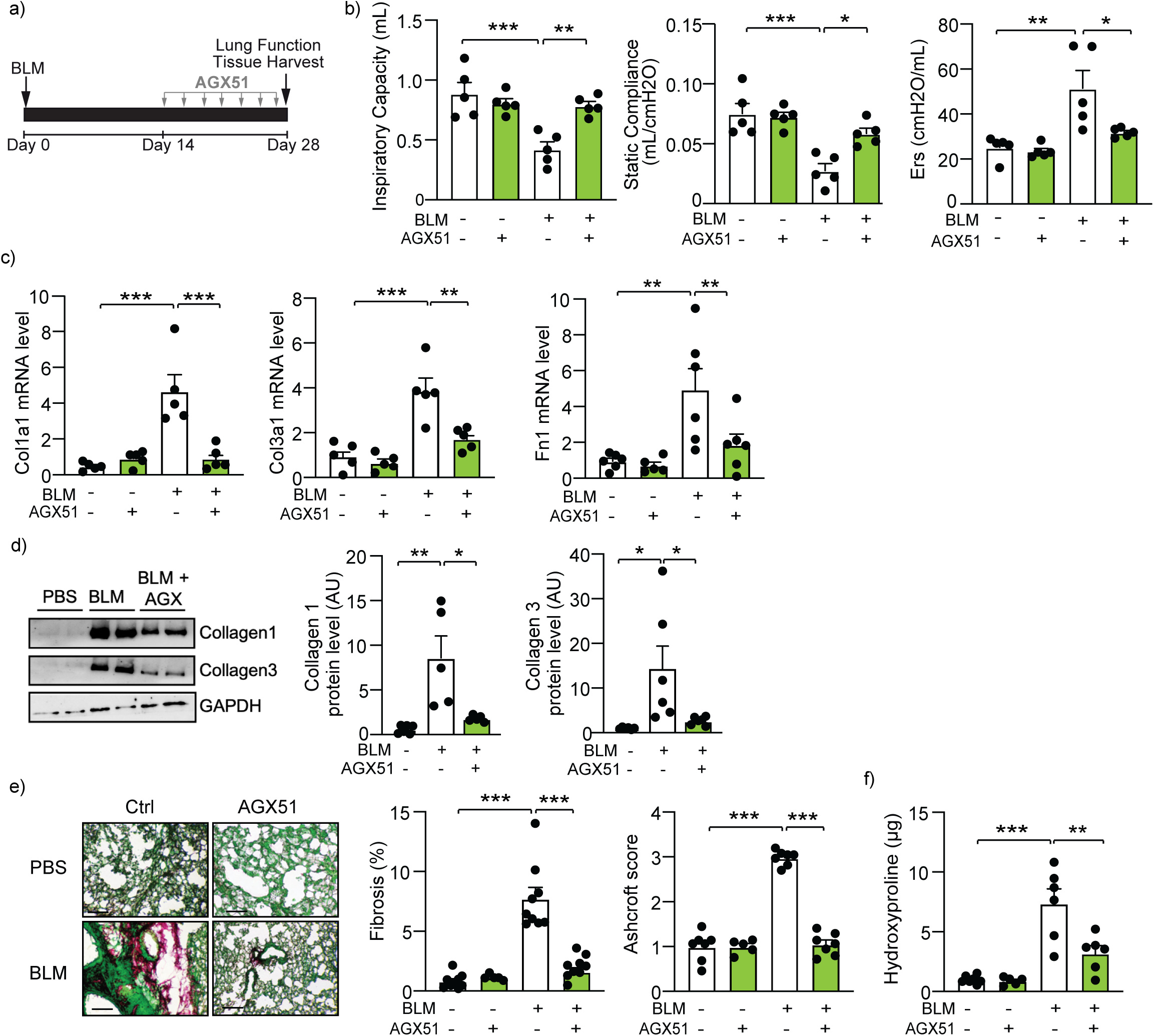
A pharmacological inhibition of ID1/ID3 protects mice from pulmonary fibrosis. a) Design of the study. b) Lung function data parameters for inspiratory capacity, compliance and single frequency elastase n=5 mice/group. c) PCR analysis of Col1a1, Col3a1 and Fn1 mRNA levels in lungs of the indicated groups n=5-6 mice/group. d) Lung protein expression of Collagen- I and Collagen-III in lungs from the indicated groups n=5-6 mice/group. e) (Left) Representative images from Fast Green/Sirius-Red-stained lungs of the indicated groups. n=5-9 mice/group. Scale bars: 200 μm. (Middle) Quantitative analysis of fibrosis. (Right) Ashcroft scores representing the extent of fibrosis. n = 5-7 mice/group. f) Hydroxyproline content in lungs from the indicated groups n = 5-7 mice/group.* P < 0.05; ** P < 0.01. *** P < 0.001.

### A lung specific inhibition of ID1/ID3 attenuates pulmonary fibrosis

To manipulate the expression of ID1 and ID3 specifically in the lung, we generated Adeno-Associated Viruses 1 (AAV1) expressing short hairpins targeting ID1 and ID3 (shID1 and shID3) and delivered them via an intratracheal injection to pulmonary fibrosis-diseased mice two weeks after BLM injection (figure 6a). BLM induced a significant increase in ID1 and ID3 mRNA levels in AAV1-Ctrl-treated mice, whereas AAV1-shID1/ID3 treatment resulted in a marked decrease in ID1 and ID3 mRNA levels (figure 6b). The intratracheal delivery of AAV-shID1/ID3 did not induce any significant changes in ID1 and ID3 mRNA levels in the heart, liver and kidney (supplementary figure S5). Mice that received an intratracheal injection of AAV-shID1/ID3 did not show any phenotype under basal conditions (figure 6c-f). In mice treated with BLM, AAV- shID1/ID3 delivery improved the lung function (i.e., increased inspiratory capacity, increased static compliance, and decreased respiratory elastance; figure 6c). AAV-shID1/ID3-treated mice exhibited a significant reduction in Col1a1, Col3a1 and Fn1 mRNA levels (figure 6d). Additionally, lungs of AAV-shID1/ID3-treated mice displayed a significant decrease in hydroxyproline level (figure 6e). Histological analysis of lung tissues with Sirius Red/Fast Green staining revealed a fibrotic response in AAV-Ctrl-treated mice, whereas AAV-shID1/ID3 resulted in a significant decrease of lung fibrosis (figure 6f). Consistently, AAV-shID1/ID3-treated mice presented with a lower Ashcroft score when compared to AAV-Ctrl-treated mice (figure 6f). These results indicate that a lung specific inhibition of ID1 and ID3 protects mice from lung fibrosis.

**Figure 6:**
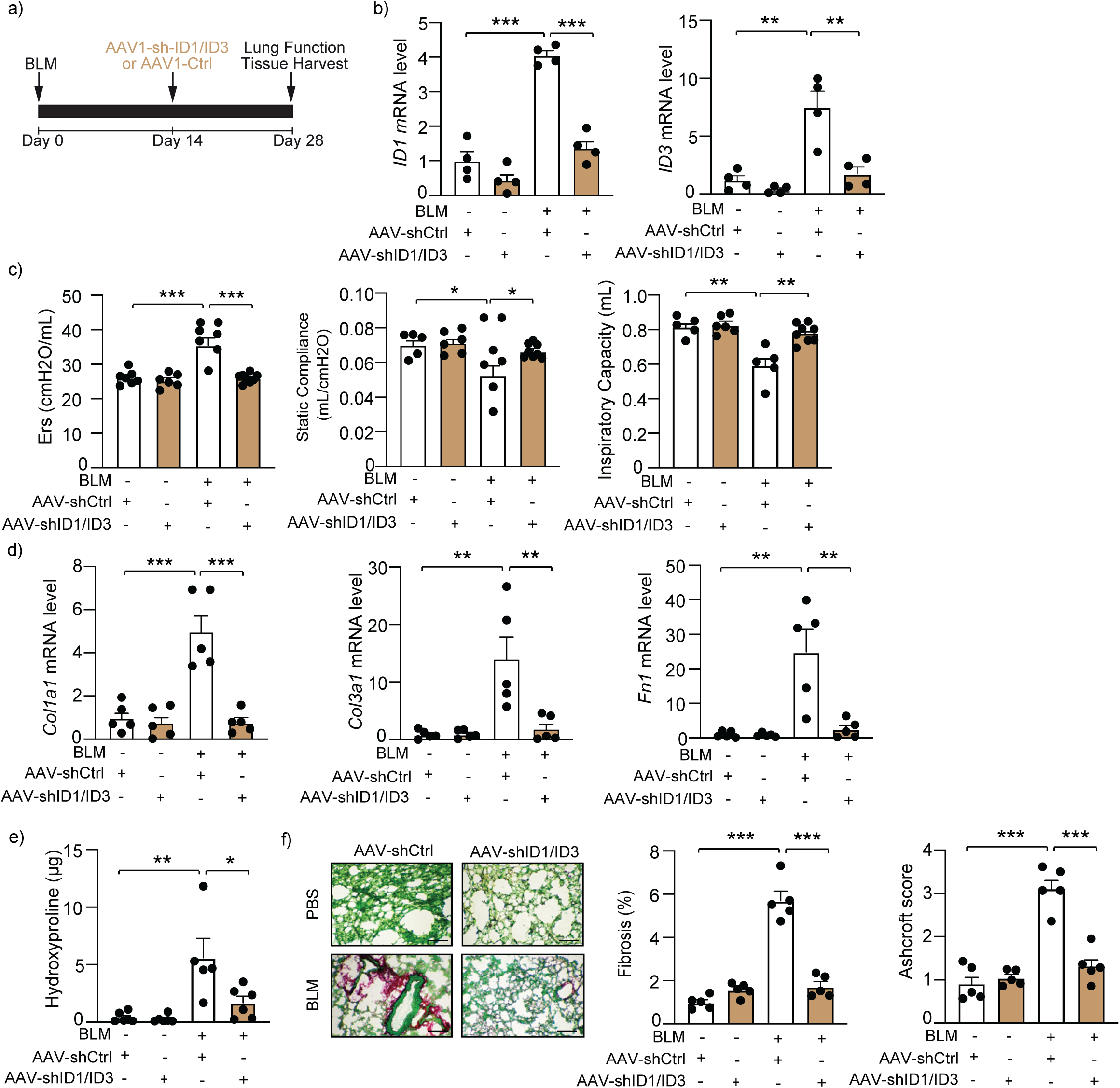
A lung specific inhibition of ID1 and ID3 decreases Bleomycin-induced lung fibrosis. a) A schematic diagram showing design of the study. b) PCR analysis of ID1 and ID3 mRNA levels in lungs of the indicated groups n=4 mice/group . c) Lung function data parameters for inspiratory capacity, compliance, and single frequency elastase n=5-8 mice/group. d) Col1a1, Col3a1 and Fn1 mRNA expression in lungs from the indicated groups n=5 mice/group. e) Hydroxyproline content in lungs from the indicated groups n=5-6 mice/group. f) (Left) Representative images from Fast Green/Sirius-Red-stained lungs of the indicated groups n=5- mice/group. Scale bars: 200 μm. (Middle) Quantitative analysis of fibrosis. (Right) Ashcroft scores representing the extent of fibrosis. n = 5 mice/group. * P < 0.05; ** P < 0.01. *** P < 0.001.

### ID and ID3 control cell cycle genes and pMEK1 signaling pathways

By analyzing the binding sites of transcription factors on the promoters’ regions of ID1 and ID3 using PROMO v3.0.2 database (15), we found Egr-1 predicted to bind in ID1 and ID3 promoters’ regions (16). Importantly, the expression level of Egr-1 is known to be increased in IPF (17), and Egr-1 is known to be involved in pulmonary fibrosis (18). Egr-1 overexpression induced a significant increase in ID1 and ID3 mRNA (supplementary figure S6a) and protein (supplementary figure S6b) levels. Given that hypoxia has long been implicated in the pathogenesis of fibrotic diseases and that several studies have demonstrated a direct link between hypoxia and the development of IPF (19, 20), we investigated whether hypoxia regulates ID1 and ID3 levels. Hypoxia (1% O2) induced a significant increase in ID1 and ID3 mRNA levels (supplementary figure S6c).

To elucidate the functional role of ID1 and ID3 in pulmonary fibrosis, it is critical to identify their mechanism(s) of action. To this purpose we performed RNA sequencing analysis on 4 biological replicates per group of RNA extracted from human lung fibroblasts treated with a specific inhibitor of ID1 and ID3 (AGX51) or PBS in the presence of serum (5% FBS). Among the regulated mRNAs, 369 downregulated and 483 upregulated genes were differentially expressed upon ID1/ID3 inhibition. The functional categories of genes significantly enriched upon ID1/ID3 inhibition using KEGG pathway revealed that the majority of regulated pathways in HLFs treated with AGX51 were associated with the cell cycle (figure 7a). We found cyclin A2 (Ccna2), cyclin B2 (Ccnb2) and cyclin-dependent kinase 1 (Cdk1) among the regulated genes. To validate the RNA seq results, we isolated mRNA from HLFs that had been treated with AGX51 and subjected it to qPCR. AGX51 reduced Ccna2, Ccnb2 and Cdk1 mRNA levels (figure 7b). BLM treatment induced a marked upregulation of pulmonary Ccna2, Ccnb2 and Cdk1 mRNA levels. This upregulation was significantly reversed by ID1/ID3 pharmacological inhibition (figure 7c). Validation of the qPCR data by immunoblotting in cells stimulated with serum and lungs of BLM-treated mice confirmed higher amounts of Ccna2, Ccnb2 and Cdk1; whereas ID1/ID3 inhibition decreased their protein levels (figure 7d-e).

**Figure 7:**
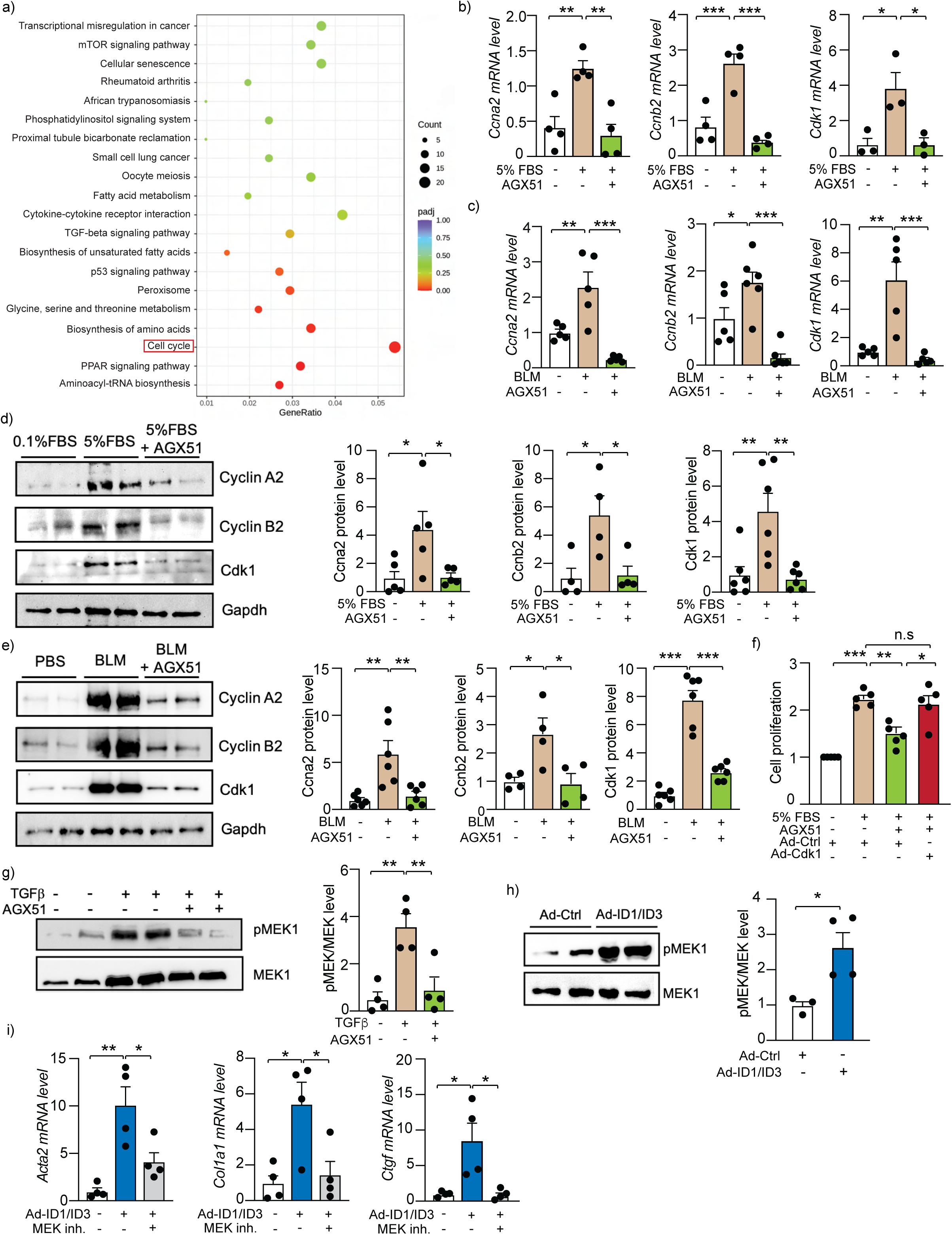
ID1/ID3 act through cell cycle genes and the MEK pathway. a) Scatter plot of top KEGG pathways enriched by differentially expressed genes in PBS-treated HLF vs AGX-51- treated HLF in the presence of Serum (5% FBS) (n = 4 per group). b) PCR analysis of ccna2, ccnb2 and cdk1 mRNA levels in HLF treated with PBS or AGX51 in the presence of Serum (5% FBS) (n = 3-4 per group). c) Pulmonary ccna2, ccnb2 and cdk1 mRNA levels in lungs from the indicated groups (n = 5-6 per group). d) CCNA2, CCNB2 and CDK1 protein levels in HLF treated with 5%FBS in the presence or absence of AGX51 (n = 4-6 per group). e) Lung protein expression of CCNA2, CCNB2 and CDK1 in lungs from the indicated groups (n = 4-6 per group). f) Proliferation of human lung fibroblasts in the presence of the indicated treatments. n = 5 experiments performed in triplicate. g) HLF treated with TGFβ or AGX51 were lysed and analyzed on western blots with antibodies against MEK1 and phospho-MEK1. (Left) Representative Western blots. (Right) Quantitative analysis of Western blot data (n = 4 per group). h) HLF treated with Adenovirus control (Ad-Ctrl) or Adenoviruses ID1+ID3 (Ad-ID1/ID3) were analyzed on western blots with antibodies against MEK1 and phospho-MEK1. (Left) Representative Western blots. (Right) Quantitative analysis of Western blot data (n = 3-4 per group). i) qPCR assessment of Acta2, Col1a1 and CTGF mRNA levels 48h after HLF treatment with Adenoviruses ID1+ID3 (Ad-ID1/ID3) and a MEK1 inhibitor (Selumetinib, 10 µM). n = 4 experiments performed in duplicate. * P < 0.05; ** P < 0.01. *** P < 0.001.

We next conducted a rescue experiment to validate the functional relevance of the cell cycle genes pathway in mediating ID1/ID3 effect. HLF were treated with AGX51 in the presence of an adenovirus encoding Cdk1 or an adenovirus control. ID1/ID3 inhibition induced a decrease in HLF proliferation, yet cdk1 overexpression largely abrogated the anti-proliferative effect of AGX51 (figure 7f).

Given that ID1 and ID3 are known to regulate the MEK/ERK pathway in cancer (7) and that MEK is known to regulate lung fibroblast differentiation into myofibroblast (21), we next assessed the effect of ID1/ID3 inhibition on MEK1 phosphorylation in human lung fibroblasts. Western-blot analysis revealed that ID1/ID3 inhibition decreases MEK1 phosphorylation in NHLF treated with TGFβ1 (figure 7g), whereas ID1/ID3 overexpression increases p-MEK1 level in human lung fibroblasts (figure 7h). Elevation of ID1/ID3 levels induced fibroblasts differentiation into myofibroblasts (determined by increased Acta2, Col1a1 and Ctgf mRNA levels), yet MEK1 inhibition abrogated the effect of ID1/ID3 (figure 7i). These results indicate that ID1/ID3 act on lung fibroblast proliferation through the Ccna2/Ccnb2/Cdk1 pathway and inhibit fibroblast differentiation into myofibroblast through the MEK/ERK pathway.

## Discussion

Ongoing research aims to better understand the pathophysiology of IPF and define a new therapeutic approach to hinder this devastating disease. As approved therapies are not sufficient to fully halt the disease progression of IPF, there is significant unmet medical need. In this study, we focused on the role of ID proteins and their effects on pulmonary fibrosis, as well as their potential as a therapeutic target. ID proteins are a family of highly conserved transcriptional regulators that play a pivotal role during the developmental processes (22). We found ID1 and ID3 mRNA levels to be significantly elevated in lungs and lung fibroblasts isolated from human patients and mice with pulmonary fibrosis. Additionally, we found ID1 and ID3 levels to be increased in human lung fibroblasts following TGF-β1 treatment. We demonstrated that ID1/ID3 simultaneous inhibition decreases lung fibroblast proliferation, migration and differentiation into myofibroblast. We further showed that genetic and pharmacological inhibition of ID1/ID3 improve lung function and protect mice from the development of pulmonary fibrosis. Mechanistically, we found that ID1/ID3 regulate cell cycle genes and the MEK pathway.

ID proteins are primarily expressed during development and are not typically found in adult tissues, nonetheless they have been reported to be re-expressed in numerous human cancers (23). Increased expression of ID proteins, especially ID1, ID2, and ID3 has been associated with advanced tumor stage and poor prognosis in many types of human cancers (24). ID1 level was shown to be increased in the lungs of BLM-treated mice (25, 26); however, ID1 regulation in human IPF and ID3 expression in the lungs of patients and mice with pulmonary fibrosis has never been investigated. Importantly, we established in this study that ID1 and ID3 levels are increased in lungs fibroblasts of patients with IPF, in healthy lung fibroblasts treated with TGF-β, and in lungs and lung fibroblasts of mice with pulmonary fibrosis. These findings are of great relevance when analyzing the IPF disease.

ID gene expression and protein abundance are known to be regulated by a wide range of growth factor and cytokine signaling cascades, as well as by post-translational modifications and degradation (27). Our results indicate that ID1 and ID3 levels are increased upon hypoxia exposure. Hypoxia regulates the expression of many genes through hypoxia-inducible factors (HIFs) (28). HIF-1 and HIF-2 are involved in the proliferation, differentiation, extracellular matrix deposition, and alteration in the cell cycle of lung fibroblasts (29, 30). It is still unclear whether hypoxia exposure regulates ID1 and ID3 levels through HIF-1 or HIF-2.

The development of pulmonary fibrosis is generally preceeded by lung inflammation not resolved over time. The lung inflammation can be developed by radiation, chemotherapy, air polluants or bacterial/viral infections (31, 32). This study indicates that ID1 and ID3 levels are increased in human lung fibroblasts treated with TGF-β, a secreted cytokine that plays an important role in IPF. However, it is still unclear whether ID1 and ID3 levels are regulated by other cytokines released during lung inflammation (e.g. TNFα, IL-1, IL-6) and whether the inhibition of ID1/ID3 affects lung inflammation.

Mechanistically, we found that ID1/ID3 target key pathways involved in fibroblast proliferation, migration and differentiation into myofibroblast. ID1/ID3 inhibition reduced cyclin A2, cyclin B2 and Cdk1 mRNA levels. Importantly, Cdk1 overexpression prevented the anti-proliferative effect of ID1/ID3 inhibition. Additionnaly, ID1/ID3 overexpression increased MEK1 phosphorylation in human lung fibroblasts, and MEK1 inhibition abrogated the effect of ID1/ID3 on lung fibroblast differentiation into myofibroblast.

It was reported that BLM-treated ID1 KO mice display increased lung collagen accumulation (25, 26); however, given that ID1 and ID3 have overlapping and synergistic functions (33, 34) and that ID1 and ID3 are known to compensate for the loss of each other ((35–38) and figure 2a), a simultaneous inhibition of ID1 and ID3 is essential for the characterization of their role in vitro and in vivo. The reported increase in collagen production in the lungs of ID1 KO mice (25, 26) is most likely due to a compensatory mechanism by ID3. This study is the first to manipulate ID1 and ID3 simultaneously to study their roles in IPF. Given that ID1/ID3 double knockout mouse embryos die at mid-gestation, we generated mice carrying a ubiquitous deletion of ID3 with mice carrying a fibroblast specific deletion of ID1. The developed mice allowed us to demonstrate that a simultaneous knockdown of ID1 and ID3 protects mice from the development of pulmonary fibrosis.

A small-molecule pan-ID antagonist (AGX51) has been previously been identified as a specific inhibitor for ID1 and ID3 (39). Approximately 350 molecules were tested for their ability to perturb ID proteins interactions with target bHLH protein E47. The top candidate from this screen, AGX51, led to destabilization of ID1 and ID3 proteins and their degradation (39, 40). In pre- clinical experiments, AGX51 phenocopied the genetic ID1/ID3 loss and induced strong anti-tumor effects (14). In our study, AGX51 reversed IPF-diseased HLF dysfunction in vitro, improved the lung function of bleomycin-treated mice and decreased lung fibrosis in PF-diseased mice. ID1/ID3 pharmacological inhibition holds promise for treating IPF. Nonetheless, to minimize the risk of unwanted side effects in other organs, ID1/ID3-directed therapy against pulmonary remodeling would benefit from targeted delivery to the lung.

To manipulate ID1 and ID3 specifically in the lung, we developed AAV1 expressing short hairpins targeting ID1 and ID3 under the control of U6 promoter and delivered them via an intratracheal injection to mice. The delivery of AAV1-shID1/ID3 improved lung function and resulted in a significant decrease of lung fibrosis. Although not inducing any significant changes in ID1 and ID3 mRNA levels in other organs, AAV1 likely infects many cell types in the lung. In future studies, we aim to specifically target ID1 and ID3 in lung fibroblasts. An AAV1-shID1/ID3 under the control of a fibroblast specific promoter could be used to determine the effect of a fibroblast specific-ID1/ID3 inhibition on pulmonary fibrosis.

In summary, we propose a model in which ID1 and ID3 are upregulated by TGF-β1, hypoxia and Egr-1 in lung fibroblasts. A simultaneous inhibition of ID1/ID3 decreases human lung fibroblast proliferation, migration and differentiation into myofibroblast. Our study demonstrates that a genetic or pharmacological inhibition of ID1 and ID3 improves lung function and reverses pulmonary fibrosis in mice.

## Acknowledgments

We thank Emerson Obus and Roslyn Fawbush for technical assistance. We gratefully acknowledge support of this work by the American Heart Association (24POST1189241), the DC Women’s Board and the National Institutes of Health (R01HL160963).

## Disclosures

Y.S. and S.A. have a pending patent on the inhibition of ID proteins to treat IPF. R.B own founder shares of Angiogenex, Inc. The other authors declare no conflict of interest.

## Notes

### Competing Interest Statement

The authors have declared no competing interest.

