## Supplemental Material for "A Simultaneous Inhibition of ID1 and ID3 Protects Against Pulmonary Fibrosis"

### Supplementary Methods

#### Reagents

AGX51 (MedChem Express) was used at a final concentration of 20 μM. AAV1-GFP-U6-mID1-shRNA, AAV1-GFP-U6-mID3-shRNA, Ad-h-ID1, Ad-h-EGR1, Ad-h-ID3 and Ad-Cdc2 were purchased from Vector Biosystem. Recombinant human Transforming growth factor-β 1 (TGF-β1, Peprotech) aliquoted and kept at -20 °C until use. TGF-β1 was used at concentration of 5 ng/ml. MEK1 inhibitor (Selumetinib, MedChem Express) was aliquoted and kept at -80 °C until use.

#### Generation and use of ID1/ID3 knock-out mice

ID1/ ID3 double knock-out mice were generated by crossing mice carrying a ubiquitous deletion of ID3 with mice carrying a fibroblast specific deletion of ID1 (by crossing $ID1^{fl/fl}$ mice with Col1a2-CreER mice (JAX stock #029567)). Genotyping of these mice was performed by PCR. All animal procedures were conducted according to the Institutional Animal Experimentation Guidelines. The mice were housed in a pathogen-free facility, and all animal experiments were approved by the Animal Care and Use Committees of Virginia Tech.

#### siRNAs transfection

NHLF and IPF cells were seeded at 50,000 cells/well in 12-well plates and maintained in a 37$^{o}$C incubator with 5% $CO_2$ for 24 h before transfection with the different siRNAs. siRNA-ID1 (Assay ID S7104), siRNA-ID3 (assay ID S7112) and negative control siRNA (Assay ID 4390844) were purchased from ThermoFisher. The siRNAs were transfected into the cells using lipofectamine 2000 (ThermoFisher), according to the manufacturer's instructions.

**Cell proliferation**

Fibroblast proliferation was measured by 5-bromo-2'-deoxyuridine (BrdU) incorporation. NHLF and IPF cells were seeded at 4000 cells/well in 96-well plates and maintained in a 37$^{o}$C incubator with 5% $CO_2$ for 24 hr before treatment. One day later, the cells were incubated with the different treatments for 48hr. Two days later, BrdU was added and incubated for 2-24hr. Cell Proliferation ELISA, BrdU (colorimetric) assay was performed (Sigma) according to the manufacturer's instructions.

**Cell migration**

Fibroblasts were seeded at a concentration of 50,000 cells/well in a 12 well plate and allowed to incubate until confluency. A scratch was then made in each well using a 200µl pipette tip. The media was then removed and cells were washed with PBS before receiving the different treatments. The scratches of each well were analyzed by optical microscopy 0 and 48 hr following the induced damage. The area between each edge of the scratch was measured using ImageJ software and expressed as percentage of closure of the area as compared to control untreated cells.

**Quantitative RT-PCR**

Total RNA was extracted from lung tissue samples in 1 ml TRI Reagent (Fisher Scientific) according to the manufacturer's instructions. RNA was reverse transcribed using the High capacity cDNA synthesis kit (Fisher Scientific). Real-time qPCR was conducted using the PerfeCTa SYBR$^{TM}$ Green PCR Fast Mix Kit (VWR) following the manufacturer's protocol. All data were normalized to the expression of housekeeping gene GAPDH. The primers sequences are listed in Supplementary Table S1.

**Immunoblot analysis**

To analyze the protein content, cell lysates were prepared using RIPA lysis buffer ((R0278-500ML, Sigma-Aldrich) supplemented with a protease and a phosphatase inhibitor cocktail (Sigma-Aldrich). After 30 minutes centrifugation at 12000xg, protein concentrations were determined using the bicinchoninic (BCA) assay (Sigma-Aldrich). The proteins were then separated using SDS-polyacrylamide gel electrophoresis (PAGE) and transferred onto nitrocellulose membranes. The membranes were blocked with 5% bovine serum albumin (BSA).

The membranes were then incubated overnight with the primary antibodies listed in supplementary Table S2. Next, the membranes were incubated with the corresponding HRP-conjugated secondary antibodies (Cell Signaling Technology). Finally, the luminescence signal was visualized using Pierce™ ECL Western Blotting Substrate (ThermoFischer Scientific).

**Hydroxyproline content**

Hydroxyproline was measured using the Hydroxyproline Assay Kit (Sigma-Aldrich, (MAK569-1KT) according to manufacturer's instructions. Left lung lobes were homogenized and 10 mg used for the quantification of the hydroxyproline content that was measured as an indicator of collagen accumulation, using a colorimetric assay.

**Sirius Red Fast Green staining**

Lung tissues were collected, infused with a PBS/OCT mixture (50:50), and stored in OCT at -80$^{o}$C. Sections (8 μm) were carefully sliced and affixed to Color Frost glass slides (ThermoFisher Scientific). For histological analysis, lung tissue sections were subjected to Sirius Red staining Fast Green FCF (F7252-5G, Sigma-Aldrich), Direct Red 80 (365548-5G, Sigma-Aldrich). Images were obtained using a light microscope. Image J software was used to quantify the collagen deposition. The Ashcroft score was used to assess pulmonary fibrosis using a validated semi-quantitative approach. This scoring system ranges from 0 (normal lung) to 8 (complete fibrous obliteration of the field).

**Lung function**

Anesthetized mice were intubated with a sterile blunt 19G cannula, which was then connected to a flexiVent respirator (SCIREQ). The 19G catheters were selected because they did not cause any injury to the trachea during intubation. Prior to each experiment, the flexiVent system was calibrated to account for the mechanical properties of both the system and the tracheal cannula, ensuring accurate calculations of lung mechanical properties. Ventilation was set with a tidal volume of 10 ml/kg body weight, a respiratory rate of 150 breaths per minute, and a positive end-expiratory pressure (PEEP) of 3 cm $H_2O$. Two recruitment maneuvers (deep lung inflation at 30 cm $H_2O$) and three pressure-controlled quasi-static pressure–volume (PV) loops were applied. The inspiratory capacity (IC), which indicates the amount of air that can be inhaled following a normal expiration, was calculated during deep inflation. This maneuver helps open closed lung areas, restore airway patency, and normalize lung volume. Tissue elastance (H), which reflects lung resistance to distension under mechanical load, was calculated by fitting the constant-phase model to impedance spectra obtained during FOT. Tissue elastance serves as an indicator of lung stiffness and typically increases in the presence of developing lung fibrosis. Quasi-static compliance (Cst), which represents lung distensibility, was determined as the mean of the three PV loop values using the Salazar-Knowles equation.

**Supplemental Table S1. Real-time PCR Primers for analysis**

| Gene | Direction | Primer sequence (5'-3') |
|---|---|---|
| Human α-SMA | Forward | ACCCACAATGTCCCCATCTA |
| | Reverse | GAAGGAATAGCCACGGCTCAG |
| Human Ccna2 | Forward | TCCCCAGACTTTTTCGCTCT |
| | Reverse | GGATGCCAGTCTTACTCATAGC |
| Human Ccnb2 | Forward | TACTGCTCTGCTCTTGGCTT |
| | Reverse | GCTGTTCAACATCAACCTCCC |
| Human Cdk1 | Forward | GCCCTTTAGCGCGGATCTAC |
| | Reverse | AGGAACCCCTTCCTCTTCACT |
| Human Col1a1 | Forward | GGCTCCTGCTCCTCTTAGC |
| | Reverse | TCTCTTACGCAGGTGATTGGT |
| Human CTGF | Forward | TCCCAAAATCTCCAAGCCTA |
| | Reverse | GTAATGGCAGGCACAGGTCT |
| Human/ Mouse GAPDH | Forward | CCAAGGTCATCCATGACAACTT |
| | Reverse | GTCTTCTGGGTGGCAGTGATG |
| Human ID1 | Forward | CGCATCTTGTGTCGCTGAAG |
| | Reverse | CCGATCGGTCTTGTTCTCCC |
| Human ID2 | Forward | CCCACTATTGTCAGCCTGCA |
| | Reverse | CTGCAAGGACAGGATGCTGA |
| Human ID3 | Forward | TACAGCGCGTCATCGACTAC |
| | Reverse | TGACAAGTTCCGGAGTGA |
| Human ID4 | Forward | TGAGTAGTACCGGGGAGTGGG |

| | | |
|---|---|---|
| | Reverse | AAGAAAAGTAGCCCACCCGG |
| Mouse Ccna2 | Forward | CTCACTACATAGCTGACTTGGAC |
| | Reverse | CTGGGTGGCGCCTTTAATC |
| Mouse Ccnb2 | Reverse | GTTCTGAGGTTTCTTCGCCA |
| | Forward | ATCCGGCGGGCAGTTTTAG |
| Mouse Cdk1 | Forward | CTCCTGGGCAGTTCATGGAT |
| | Reverse | CCACAGCGTCACTACCTCG |
| Mouse Col1a1 | Forward | CTGGCAAGAAGGGAGATGA |
| | Reverse | CACCATCCAAACCACTGAAA |
| Mouse Col3a1 | Forward | ACAGCAAATTCACTTACACAGTTC |
| | Reverse | CTCATTGCCTTGCGTGTTT |
| Mouse CTGF | Forward | AGAACTGTGTACGGAGCGTG |
| | Reverse | CCATCTTTGGCAGTGCACAC |
| Mouse Fibronectin | Forward | ATGAGAAGCCTGGATCCCCT |
| | Reverse: | CAGTTGGGGAAGCTCATCTGT |
| Mouse ID1 | Forward | ATCATGAAGGTCGCCAGTGG |
| | Reverse | CCGACAGACCAAGTACCACC |
| Mouse ID2 | Forward | ATCCTGTCCTTGCAGGCATCT |
| | Reverse | GTCCATTCAACGTGTTCTCCTG |
| Mouse ID3 | Forward | TTTCCGAGAATGGGGTGTCG |
| | Reverse | ATCAGGGCAGCAGAGCTTTT |
| Mouse ID4 | Forward | GCAGGGTGACAGCATTCTCT |
| | Reverse | AATTTCTCCTCTGGCCCTCC |

**Supplemental Table S2. Antibodies used.**

| Antibody | Company | Product number |
|---|---|---|
| Monoclonal Anti-Actin, α-Smooth Muscle clone 1A4 | Sigma Aldrich | A2547 |
| Anti-mouse IgG, HRP-linked Antibody | Cell Signaling Technology | 7076S |
| Anti-rabbit IgG, HRP-linked Antibody | Cell Signaling Technology | 7074S |
| Anti-Cyclin A2 antibody | Abcam | EPR17351 |
| Cdc2 (E1Z6R) Rabbit mAb | Cell Signaling Technology | 28439 |
| Collagen 3 Antibody (B-10) | Santa Cruz Biotechnology | sc-271249 |
| Collagen 1 Antibody (3G3) | Santa Cruz Biotechnology | sc-293182 |
| Cyclin B2 Monoclonal Antibody (X29.2) | ThermoFisher | MA1-156 |
| Fibronectin polyclonal antibody | ProteinTech | 15613-1-AP |
| GAPDH Monoclonal antibody | Proteintech | 60004-1-Ig |
| ID1 Antibody (F-10) | Santa Cruz Biotechnology | sc-365654 |
| ID3 monoclonal antibody (2B11) | ThermoFisher | MA1-23242 |
| MEK1/2 (D1A5) Rabbit mAb | Cell Signaling Technology | 8727T |
| Phospho-MEK1/2 (Ser217/221) (41G9) | Cell Signaling Technology | 9154T |

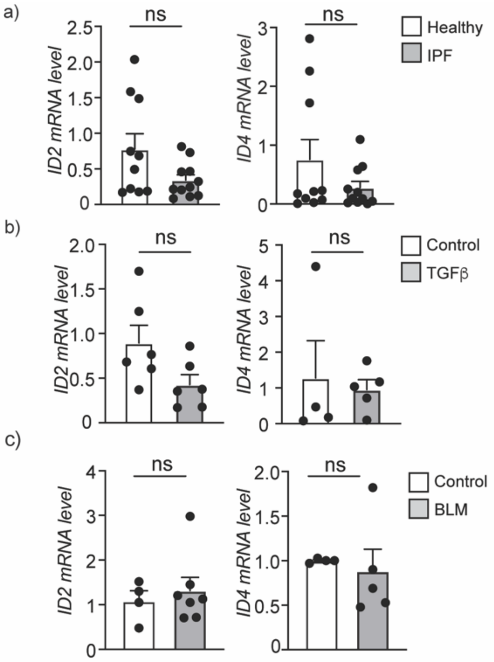


**Supplementary Figure S1: ID2 and ID4 regulation in IPF.** a) ID2 and ID4 mRNA levels determined by real time qPCR analyses in lung fibrobalsts isolated from healthy donors and patients with IPF (n=10-11/group). b) ID2 and ID4 mRNA levels in healthy human lung fibroblasts treated with TGFβ (5 ng/ml) for 48 hours. n = 5-6 experiments performed in duplicate. c) Lung expression of ID2 and ID4 mRNA levels in lung homogenates from control and BLM-treated mice (n = 4-7 mice/group). ns: not significant.

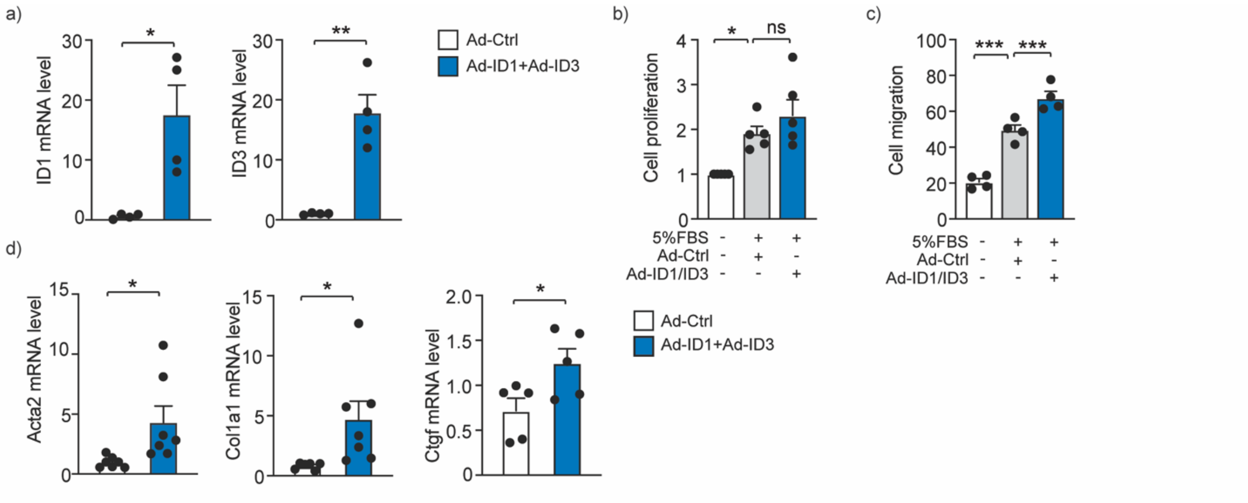


**Supplementary Figure S2: Overexpression of ID1/ID3 increases fibroblast to myofibroblast differentiation.** a) ID1 and ID3 mRNA levels in healthy human lung fibroblasts infected with Ad-Ctrl or Ad-ID1+Ad-ID3 for 48 hours. n =4 experiments performed in duplicate. b) Proliferation of healthy human lung fibroblasts infected with Ad-Ctrl or Ad-ID1+Ad-ID3 in the presence or absence of Serum (5% FBS). n = 5 experiments performed in triplicate. c) Migration of healthy human lung fibroblasts in the presence of the indicated treatments. n = 4 experiments performed in duplicate. d) Acta2, Col1a1 and Ctgf mRNA levels in healthy human lung fibroblasts infected with Ad-Ctrl or Ad-ID1+Ad-ID3 for 48 hours. n = 5-7 experiments performed in duplicate.* $P < 0.05$; ** $P < 0.01$.

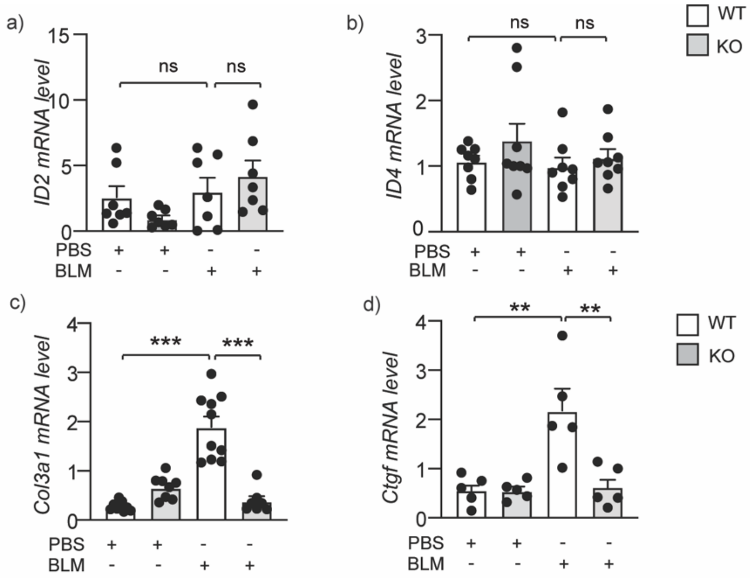


**Supplementary Figure S3: ID1/ID3 deletion decreases Bleomycin-induced lung fibrosis.** (a-b) PCR analysis of ID2 (a) and ID4 (b) mRNA levels in lungs of the indicated groups (n=7-8 mice/group). (c-d) PCR analysis of Col3a1 (c) and Ctgf (d) mRNA levels in lungs of the indicated groups (n = 7-10 mice/group in (c) and n= 5 mice/group in (d)). * $P < 0.05$; ** $P < 0.01$. *** $P < 0.001$.

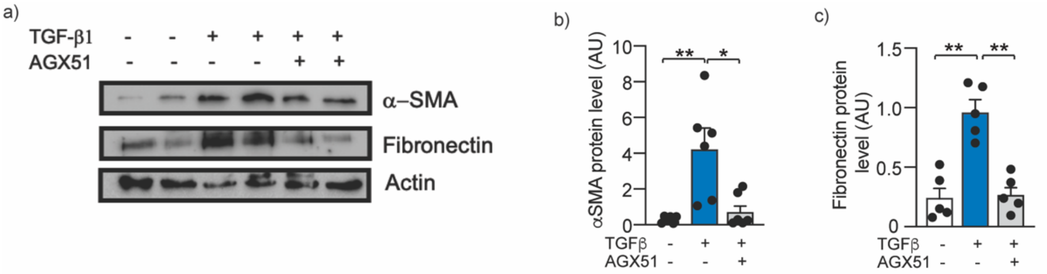


**Supplementary Figure S4: Pharmacological inhibition of ID1/ID3 decreases fibroblast differentiation into myofibroblast.** a) Western immunoblotting was performed to evaluate Fibronectin and α-SMA expression in human lung fibroblasts in the presence or absence of TGF-β1 and AGX51. b) Quantitative analysis of α-SMA protein expression. Protein expression was normalized to Actin. c) Quantitative analysis of Fibronectin protein expression. Protein expression was normalized to Actin. n = 5-6 /group. * $P < 0.05$; ** $P < 0.01$.

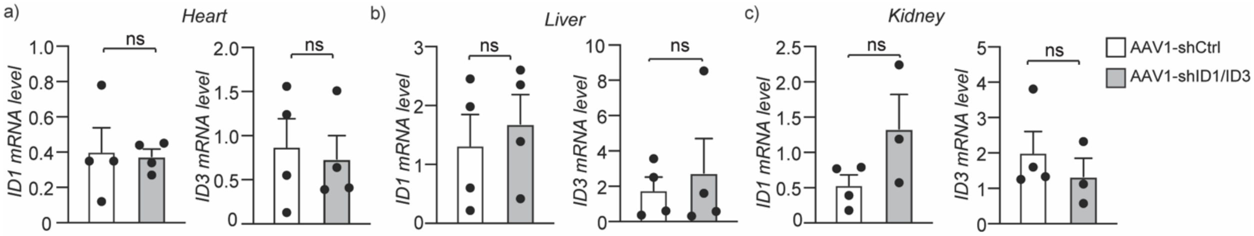


**Supplementary Figure S5: A lung specific manipulation of ID1 and ID3.** PCR analysis of ID1 and ID3 mRNA levels in (a) heart, (b) liver and (c) kidney of the indicated groups. n = 3-4 mice/group. ns: not significant.

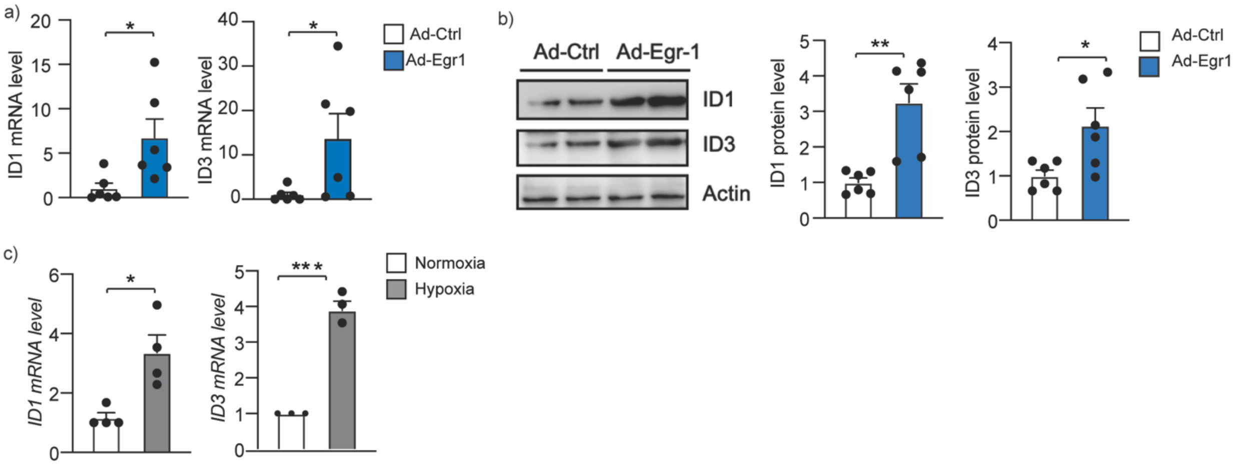


**Supplementary Figure S6: ID1/ID3 regulation by Egr-1 and hypoxia.** (a) ID1 and ID3 mRNA levels in human lung fibroblasts infected with Ad-Ctrl or Ad-Egr-1. n = 6 experiments performed in duplicate.(b) ID1 and ID3 protein levels in C2C12 cells infected with Ad-Ctrl or Ad-Egr-1. n = 6 experiments performed in duplicate. (c) ID1 and ID3 mRNA levels in human lung fibroblasts exposed to normoxic (21% $O_2$) or hypoxic (1% $O_2$) conditions for 48 hours. n = 3-4 experiments performed in duplicate. * $P < 0.05$; ** $P < 0.01$.
